# Application of Spectral Domain Optical Coherence Tomography To Guide Cochlear Implant Electrode Array

**DOI:** 10.1101/2025.05.27.656464

**Authors:** Pawina Jiramongkolchai, Marcello M. Amaral, Ratul Paul, Abby Matt, Meitong Nie, Senyue Hao, Aaron J Adkins, Hongwu Liang, Timothy Holden, Craig A. Buchman, Chao Zhou

**Affiliations:** Department of Otolaryngology – Head and Neck Surgery, Washington University in St. Louis School of Medicine - St. Louis, MO; Department of Biomedical Engineering, Washington University in Saint Louis, Saint Louis, MO; Department of Biomedical Engineering, Universidade Brasil, San Paulo, Brazil

## Abstract

**Hypothesis:** A custom spectral domain optical coherence tomography (SD-OCT) platform can be used for real-time guidance of a cochlear implant electrode array (EA).

**Background:** With current cochlear implant surgical techniques, placement of the EA is a blind maneuver in which the surgeon relies on tactile feedback as the EA advances through the cochlear lumen. Cochlear implant trauma is a leading factor for poor speech performance outcomes and loss of residual hearing following surgery. Optical coherence tomography (OCT) is a non-invasive imaging modality that provides real-time visualization of tissue microstructure at higher spatial resolutions compared to clinical CT and MRI. Already adopted as standard of care in ophthalmology, OCT has the potential to assist the surgeon in real-time visualization of the EA trajectory. Unlike commercial systems, our custom OCT system allows tailored wavelength, scanning geometry, and real-time processing, which are critical factors for navigating the compact anatomy of the facial recess to image the cochlea.

**Methods:** A custom-built SD-OCT system was used to image cochlear microanatomy in mice and human cadaveric temporal bones. The OCT system was then used to guide a mock EA in human cadaveric temporal bones in real-time using individual B scans that were reviewed sequentially as the EA was being advanced through the round window.

**Results:** Using our OCT system, high-resolution (< 5.0 μm) images of cochlear microanatomy were obtained in both mice and cadaveric human temporal bones with an image sensitivity of ∼104 dB. Following cochleostomy in cadaveric temporal bones, real-time sequential OCT B-scans were used to reliably guide placement of the EA through the scala tympani.

**Conclusion:** Our custom-built SD-OCT platform can generate high-resolution real-time visualization and orientation of mammalian cochlear microanatomy that can be used to assist with real time guidance of a CI EA. This technology has the potential to serve as a real-time surgical image guidance tool to minimize EA trauma and further our understanding of human cochlear pathophysiology.

## INTRODUCTION

Sensorineural hearing loss (SNHL) is a leading cause of disability affecting over 460 million individuals worldwide.^1^ Understanding the pathophysiology of SNHL remains, in part, limited due to the inability of current imaging modalities to provide *in vivo* high spatial resolution of the inner ear. In the United States, the development of the cochlear implant has enabled hearing restoration to over 150,000 patients with sensorineural hearing loss.^2^ There are two common surgical approaches for cochlear implantation: round window membrane insertion and cochleostomy. With either approach, the surgeon relies on tactile feedback during electrode array (EA) insertion to avoid excessive force that could cause intracochlear trauma. However, the rate of electrode array insertion trauma during cochlear implantation is as high as 30%.^3^ Insertion trauma can cause loss of any pre-existing residual hearing through multiple mechanisms: translocation of the electrode through the scala tympani into the scala vestibuli, injury to the cochlear nerve fibers, and/or reactive fibrosis and inflammation.^4-6^ Most trauma from cochlear implant electrode insertion occurs during the electrode passage within the first 180 degrees of the cochlea.^7-9^ Insertional trauma due to the absence of real-time intracochlear visualization at the time of surgery has prompted active research into the development of high-resolution intraoperative imaging to assist surgeons.

Optical coherence tomography (OCT) is a non-invasive contrast-free imaging modality that can provide high depth imaging of tissue microstructure. By measuring the backscattering of near-infrared light on biologic tissue, OCT can generate *in vivo* real-time images of tissue morphology, function, and pathology. Furthermore, the spatial resolution of OCT (<5 μm) is higher than that of conventional clinical computed tomography (CT, 0.5 mm) and magnetic resonance imaging (MRI, 1-2 mm).^10^ OCT has been adopted as standard of care in ophthalmology and has recently been used in otology to visualize the tympanic membrane and middle ear pathology *in vivo*.^11-14^ With its capability of high-resolution micron-scale imaging, there is significant research interest in the application of OCT for inner ear imaging.^15-19^

Although commercial OCT systems are available, custom-built systems enable optimization of parameters such as wavelength, scanning geometry, flexibility in operator view, and real-time processing—factors critical for imaging through the constrained anatomy of the facial recess and resolving the fine structures of the cochlea. In this study, we aim to visualize cochlear microanatomy using a custom-built spectral domain OCT platform with the goal of 1) generating high resolution images of mammalian cochlear anatomy and 2) using these high-resolution OCT images to assist with real-time visualization of EA insertion.

## MATERIAL AND METHODS

The Human Research Protection Office at the study institution deemed the cadaveric human temporal bone study exempt from Institutional Review Board review. All animal experiments were performed in compliance with guidelines approved by the Institutional Animal Care and Use Committee (IACUC) at Washington University in St. Louis.

### Customized SD-OCT Platform

Specimens in this study were imaged with a custom-built non-contact SD-OCT system. This SD-OCT platform (Figure 1) has a 5X objective lens on the sample arm and utilizes a broadband supercontinuum diode laser with a 1310-nm central wavelength and 110-nm spectral range as the light source. The interferometric signal is detected by a spectrometer on the reference arm that has a 1024-pixel, 20 kHz indium gallium arsenide (InGaAs) line-scan camera. The system can provide a transverse resolution of up to 3.5 μm and an axial resolution of up to 5 μm in tissue (refractive index = 1.33). The sensitivity of the OCT system is ∼104 dB when operating at a 20 kHz A-scan rate. OCT images were processed in a custom software, Image J, and Amira (Thermo Fisher Scientific) for volumetric renderings.

**Figure 1.**
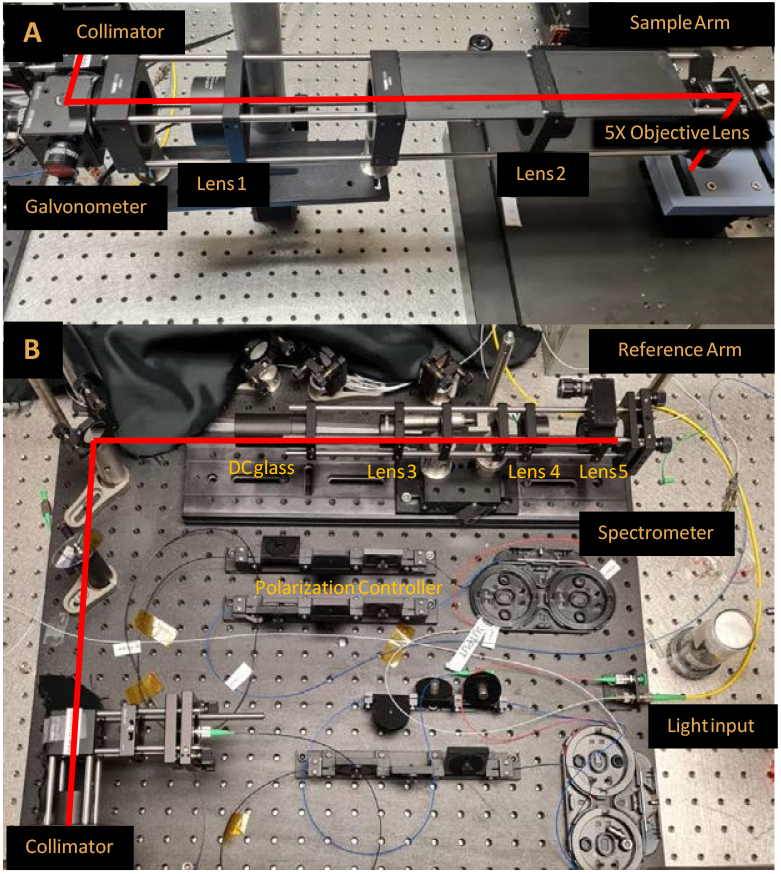
Customized 1310 nm SD-OCT platform. A) Sample arm with 5X objective lens B) Reference arm with light source and spectrometer

### Structural OCT Imaging of Mice Cochlea

Otic capsules from a total of 6 adult male wild-type C57BL/6 mice (aged ∼36 weeks) were extracted. Three of the cochleae were treated with ethylenediaminetetraacetic acid (EDTA) to decalcify the otic capsule.

### Structural OCT Imaging of Human Cadaveric Cochlea

Ten formaldehyde-fixed human cadaveric temporal bones were used and mastoidectomies performed to varying degrees to provide maximal exposure and orientation to the labyrinth. Mock cochlear implant EAs were used and inserted through a standard round window membrane (RWM) approach. OCT scans were acquired through both an intact RWM and surgically removed RWM in these temporal bone specimens.

## RESULTS

With our custom SD-OCT system, we can acquire structural imaging of mammalian cochlear microanatomy with high precision and resolution. Furthermore, we are able to use our customized SD-OCT system to guide EA insertion at the round window.

### Structural OCT Imaging of Mice Cochlea

In this *ex vivo* mouse cochlea shown in Figure 2a, the three scalae, modiolus, and Organ of Corti are visualized (Figure 2b and 2c). Figure 3 demonstrates OCT imaging of an extracted mice cochlea treated with EDTA, which improved the optical resolution and depth of penetration using our SD-OCT system. Compared to untreated cochlea, the spiral limbus, Reissner’s membrane, and Organ of Corti can be well visualized in EDTA-treated cochlea.

**Figure 2.**
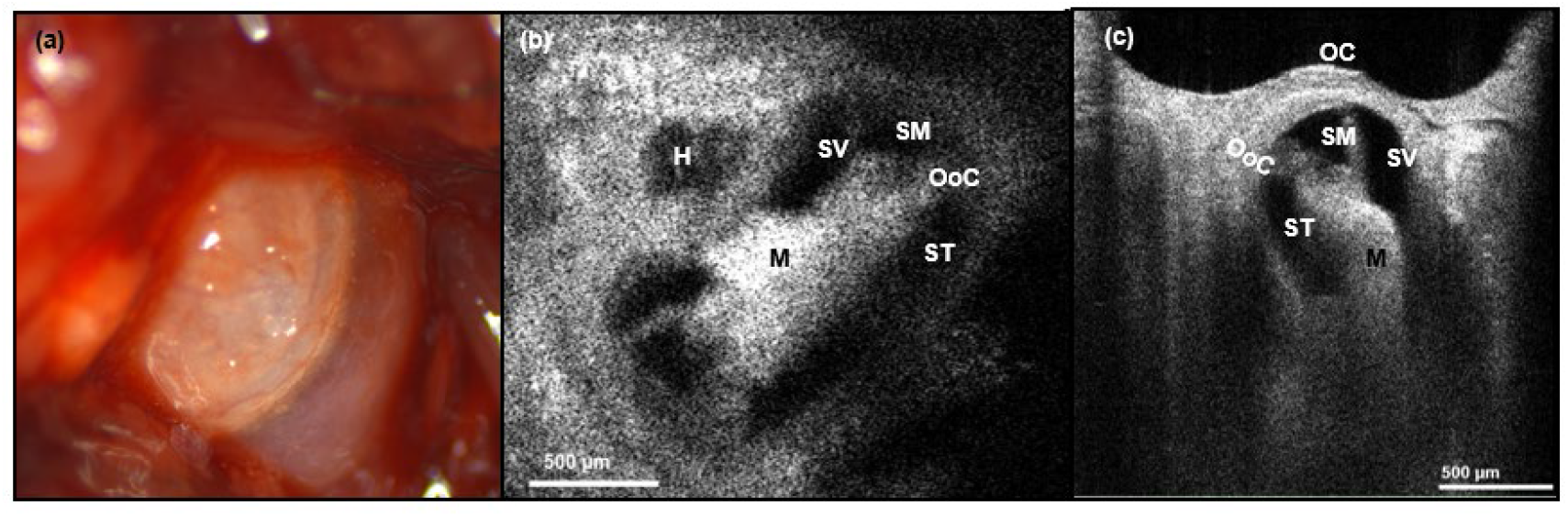
In vivo OCT imaging of a mouse cochlea. (a) Stereomicroscope of a mouse cochlea (b) 2D OCT enface view of the same cochlea specimen (c) 2D OCT orthogonal view of the cochlea. Legend: H-helicotrema, SV-scala vestibuli, SM-scala media, OoC-Organ of Corti, ST-scala tympani, M-modiolus, OC-otic capsule.

**Figure 3.**
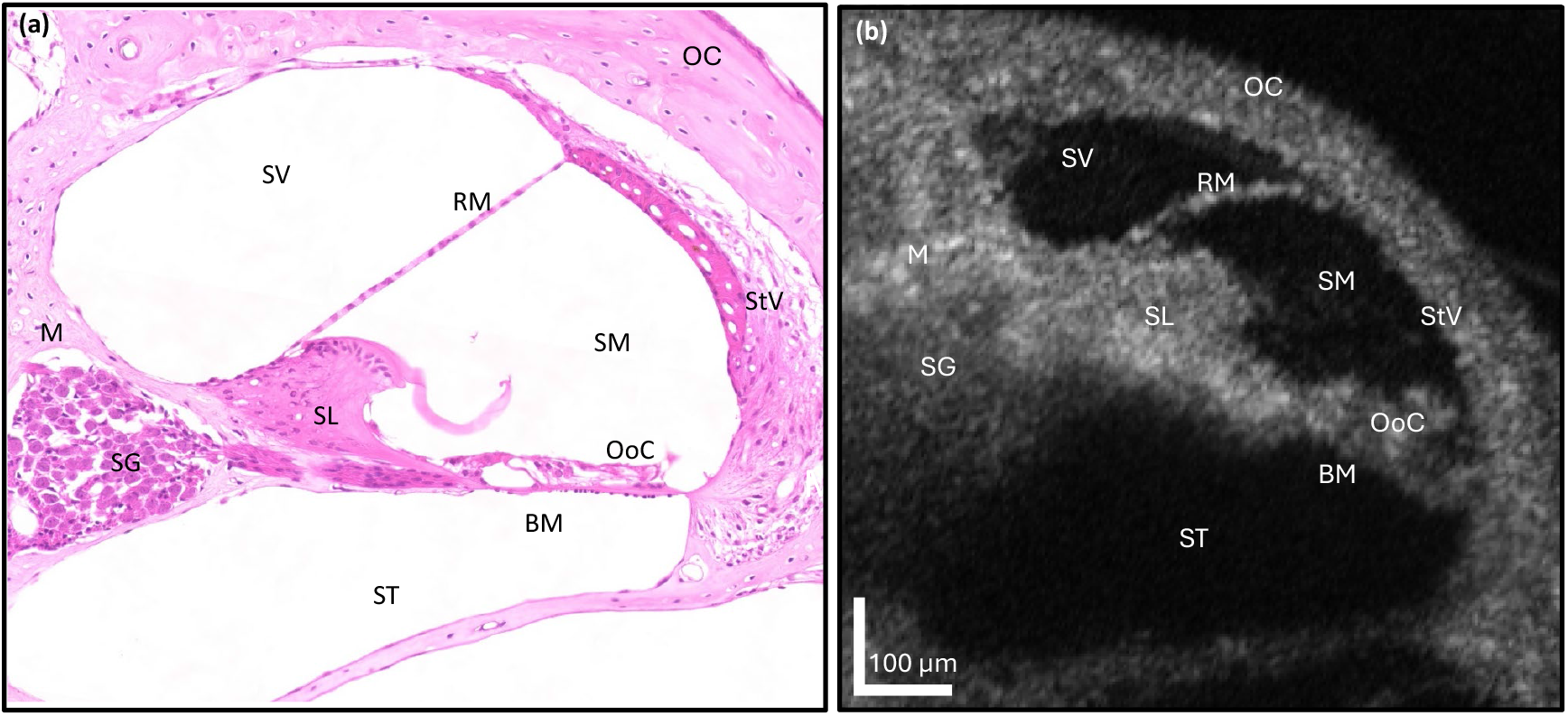
Ex vivo OCT imaging of a mouse cochlea with comparison to hematoxylin and eosin (H&E) histology. (a) H&E of the mouse cochlea (b) 2D OCT orthogonal view of the mouse cochlea. Legend: OC-otic capsule, SV-scala vestibuli, RM-Reissner’s membrane, SM-scala media, StV-strial vascularis, SL-spiral limbus, OoC-Organ of Corti, BM-basilar membrane, ST-scala tympani, M-modiolus, SG-spiral ganglion.

### Structural OCT Imaging of Human Cadaveric Cochlea

In this left-sided human cadaveric temporal bone shown in Figure 4, the vestibule was opened, and the round window membrane bony overhang drilled away to maximize exposure of the round window membrane. Because of the reduced field of view of OCT, the stereomicroscope was used in tandem with OCT to maintain anatomical spatial orientation (Figure 4A). Figure 4B is a 2D OCT cross-sectional image (B-scan) of Figure 4A with the green line indicating the horizontal cross section (4C) and the yellow line indicating the vertical cross section (4D). In Figures 4C and 4D, the scala tympani and basilar membrane can clearly be seen through an intact round window membrane. The inset of Figure 4D shows a micro-CT of a human cochlea from our institution’s micro-CT library which was used to correlate the anatomical landmarks of the round window membrane and scala tympani. Unlike the micro-CT, the basilar membrane is clearly visualized with OCT.

**Figure 4.**
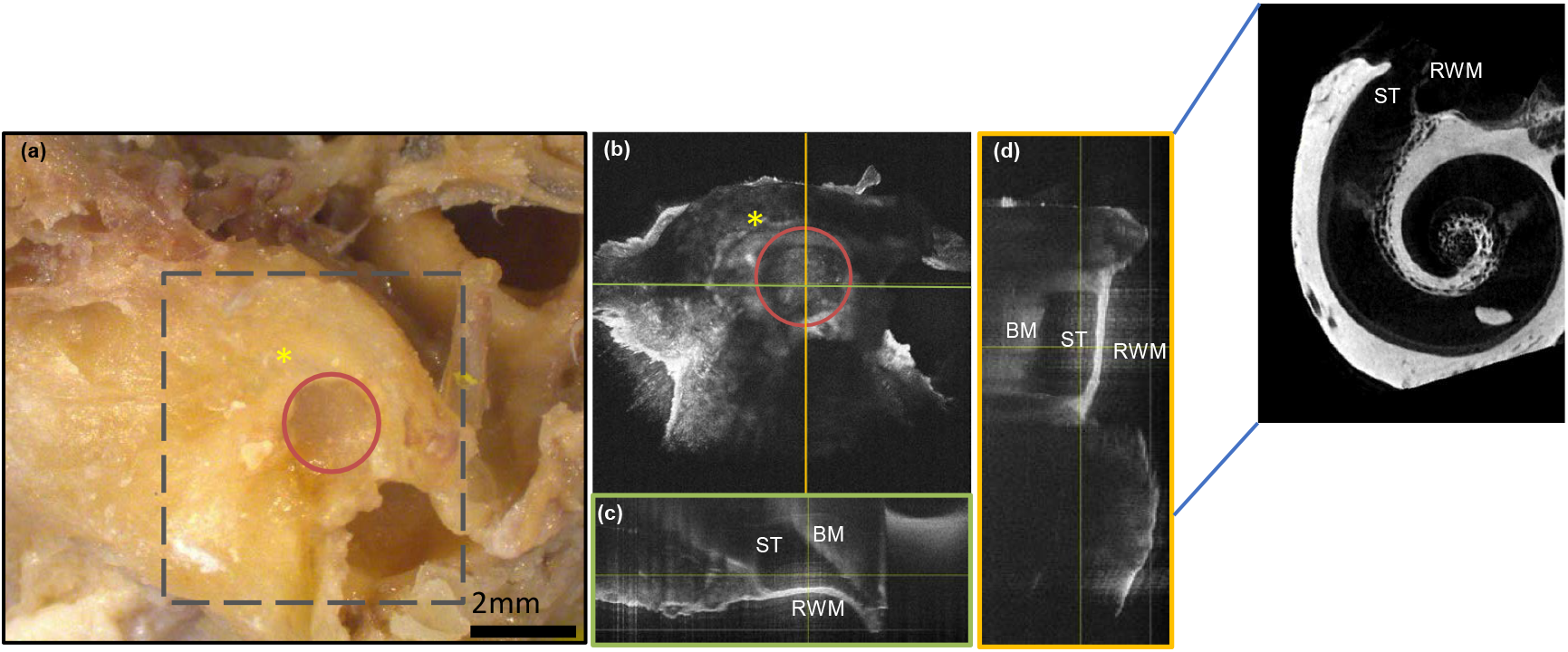
OCT imaging of a human cadaveric temporal bone through an intact round window membrane (a) steromicroscopic view (b) 2D enface OCT image with the green line indicating the horizontal cross section (c) and the yellow line showing the location of the vertical cross section (d). Legend: Dotted black square field representing field of view captured by OCT. Round window membrane (RWM, red circle); cochlear promontory (yellow asterisk); ST-scala tympani; BM-basilar membrane

### OCT Guided EA placement

The round window membrane was then surgically removed and intracochlear OCT images were obtained prior to EA placement (Figures 5A-D). With the removal of the round window membrane, the scala tympani and trajectory of the basilar membrane can be more clearly visualized. Figure 5B is a 2D OCT image of 5A from which orthogonal cross sections (Figures 5C and 5D) are obtained. Using real time orthogonal B scans (Figures 5E-G), the EA was advanced through the round window into the scala tympani. Compared to the current microscopic view during surgery, using OCT, the trajectory of the basilar membrane can be seen as the EA is being inserted, which allows the angle of insertion of the EA to be adjusted in real time to prevent basilar membrane trauma and potential translocation. Due to the high backscatter of the metal EA, only the distal tip of the electrode was visible. However, with real-time B scans, feedback about the initial trajectory of the EA was obtained enabling real-time adjustment. A flexible micro-endoscope was also used to visualize the cochlea and further confirm correct anatomic orientation with images acquired with the OCT (Figure 6).

**Figure 5.**
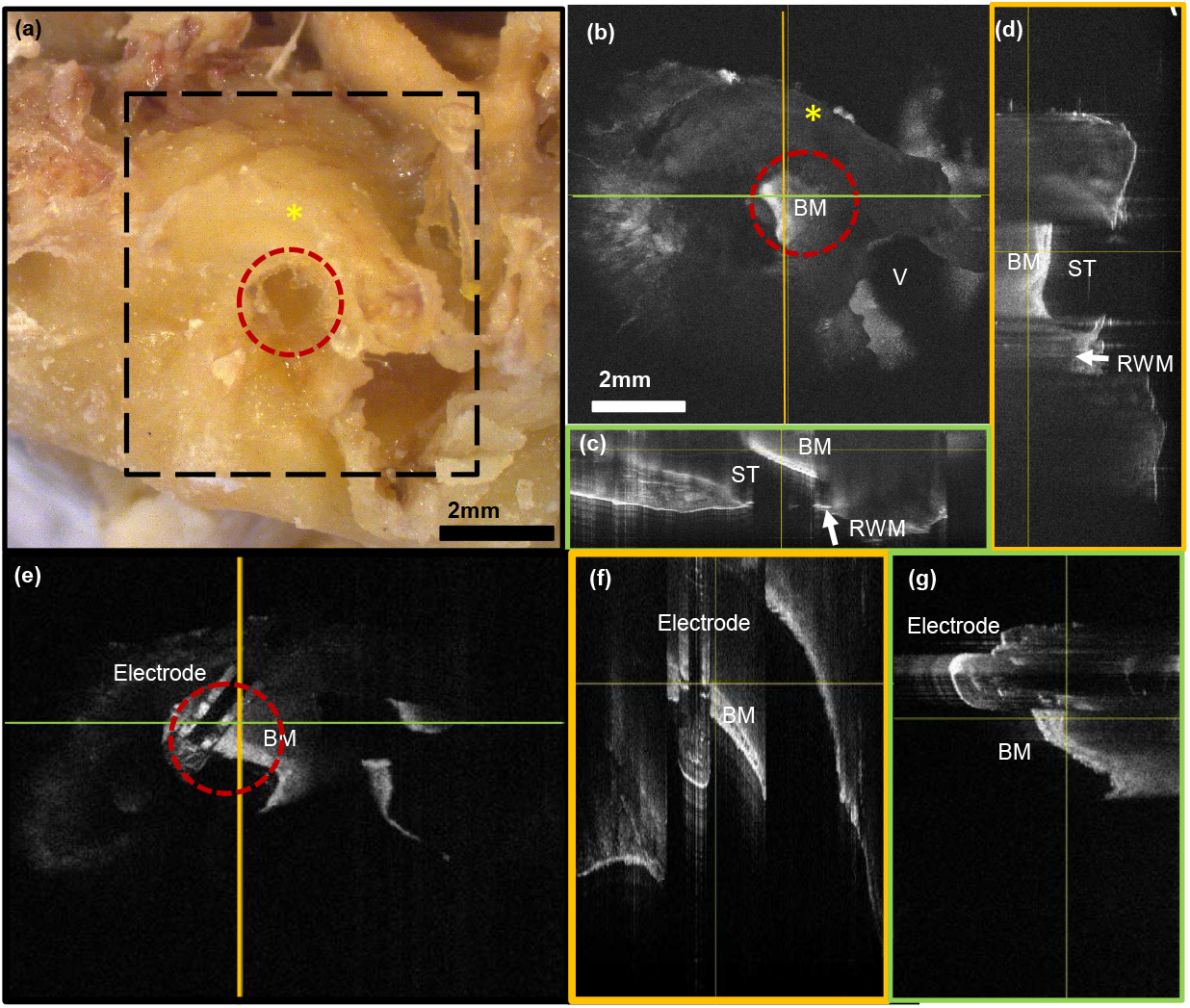
OCT images of left cadaveric human temporal bone post round window membrane removal with EA insertion. (a) Stereomicroscope image of left human cadaveric temporal bone with round window membrane nearly completely removed (dotted red line) (b) 2D enface OCT image with the green line indicating the horizontal cross section (c) and the yellow line showing the location of the vertical cross section (d). (e) 2D enface OCT image with the EA being inserted with the yellow line showing the vertical cross section (f) and the green line indicating the horizontal cross section (g). Legend: Dotted black square field representing field of view captured by OCT. Outline of the round window membrane which has been nearly completely removed (RWM, red dotted circle); cochlear promontory (yellow asterisk); V-vestibule; ST-scala tympani; BM-basilar membrane

**Figure 6.**
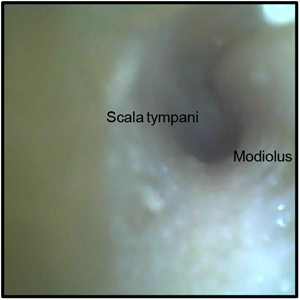
View of the cochlea including the basal turn of the scala tympani and modiolus using a flexible micro-endoscope to confirm correct anatomic orientation with images acquired with OCT.

## DISCUSSION

In this study, we have successfully demonstrated that our custom SD-OCT platform can be used for high-resolution visualization of mammalian intracochlear anatomy and for initial guidance of the EA through the round window membrane.

The use of OCT has gained increased interest due to its ability to obtain non-invasive images in real time and to potentially serve as a surgical adjunct for electrode insertion through the round window membrane. The visualization of the cochlear electrode is important to prevent traumatic injury to the delicate inner ear structures during surgery. With electrode insertion trauma, the basilar membrane, osseous spiral ligament or the spiral ligament can be damaged.^7,20,21^ In minor trauma, the electrode tip can fold over intraoperatively and in severe trauma, the electrode can migrate from the scala tympani, across the cochlear duct (i.e. scala media), and into scala vestibuli leading to complete loss of residual hearing.^22^ Anatomical variations can also affect routine insertion of the cochlear electrode. Previous studies have used OCT systems to evaluate inner ear structures of animals^23^ and human cadaveric bones.^14-16,18^

The high resolution of the images captured in this paper are 3.5 μm – 5.0 μm which is superior compared to 0.5 mm and 1-2 mm of clinically used CT and MRI, respectively^14,15^ and provide significant detail of inner ear morphology and pathology. With real-time B scans in multiple orthogonal views demonstrating the trajectory of the basilar membrane, the surgeon is able to adjust the EA trajectory at the round window. This customized OCT platform can also capture one complete volumetric dataset from a single scan in less than 30 seconds, making it potentially compatible with intraoperative use. While our images of the human cadaveric temporal bone are similar to that of other groups^16,19^, our system has a higher sensitivity (∼104 dB).

Limitations of the study include the short working distance of the SD-OCT platform and use of formaldehyde preserved cadaveric human temporal bone specimens. OCT has a known limited depth of penetration (∼1-3 mm) and due to the thickness and density of the human otic capsule, external OCT imaging is limited. As a result, the development of an OCT probe is an active area of research for us and other groups.^24^ Furthermore, although we were able to capture the Organ of Corti with our mice OCT imaging, neurosensory elements are known to rapidly degenerate in post-mortem human specimens. Future studies include the use of freshly harvested cadavers within 12-24 hours post-mortem. Finally, because OCT B-scans can present a steep surgical learning curve due to lack of spatial context, we are working on increasing the field of view with improved stitching algorithms.

Intraoperative OCT can potentially offer real-time guidance for EA insertion to minimize intracochlear trauma and provide insight into patient specific cochlear pathology. In the future, we envision an intraoperative OCT guidance system that provides visual and auditory cues, akin to car sensors, to detect intraluminal obstruction and proximity to the critical cochlear structures.

## CONCLUSION

Our customized SD-OCT platform is able to capture high-resolution images of mammalian cochlear microanatomy and provide real time guidance of EA insertion at the round window with a high sensitivity. With further refinements of stitching algorithms to improve field of view and the development of OCT probes, this technology has the potential to serve as an intraoperative guidance tool during cochlear implant surgery to minimize intracochlear trauma and further our understanding of inner ear pathophysiology.

